# High-throughput optical sensing of peri-cellular oxygen in cardiac cells: system characterization, calibration, and testing

**DOI:** 10.1101/2023.04.24.538133

**Authors:** Weizhen Li, David McLeod, John T. Ketzenberger, Grant Kowalik, Rebekah Russo, Zhenyu Li, Matthew W. Kay, Emilia Entcheva

## Abstract

Human-induced pluripotent stem cell-derived cardiomyocytes (hiPSC-CMs) represent a scalable experimental model relevant to human physiology. Oxygen consumption of hiPSC-CMs has not been studied in high-throughput (HT) format plates used in pre-clinical studies. Here, we provide comprehensive characterization and validation of a system for HT long-term optical measurements of peri-cellular oxygen in cardiac syncytia (human iPSC-CM and human cardiac fibroblasts), grown in glass-bottom 96-well plates. Laser-cut oxygen sensors having a ruthenium dye and an oxygen-insensitive reference dye were used. Ratiometric measurements (409nm excitation) reflected dynamic changes in oxygen, as validated with simultaneous Clark electrode measurements. Emission ratios (653nm vs. 510nm) were calibrated for percent oxygen using two-point calibration. Time-dependent changes in the Stern-Volmer parameter, Ksv, were observed during the initial 40 min of incubation, likely temperature-related. Effects of pH on oxygen measurements were negligible in the pH range of 4 to 8, with a small ratio reduction for pH>10. Time-dependent calibration was implemented, and light exposure time was optimized (0.6 to 0.8s) for oxygen measurements inside an incubator. Peri-cellular oxygen dropped to levels < 5% within 3 -10 hours for densely-plated hiPSC-CMs in glass-bottom 96-well plates. After the initial oxygen decrease, samples either settled to low steady-state or exhibited intermittent peri-cellular oxygen dynamics. Cardiac fibroblasts showed slower oxygen depletion and higher steady-state levels without oscillations, compared to hiPSC-CMs. Overall, the system has great utility for long-term HT monitoring of peri-cellular oxygen dynamics in vitro for tracking cellular oxygen consumption, metabolic perturbations, and characterization of the maturation of hiPSC-CMs.

## Introduction

Oxygen is important for nearly all biological processes^1^. The function of aerobic cells relying on oxidative phosphorylation, such as cardiomyocytes, is highly dependent upon oxygen availability. Precise monitoring of oxygen levels in the immediate cell vicinity (peri-cellular oxygen) will provide invaluable insight into the metabolic state of the cells. Over the last two decades, human induced pluripotent stem cell derived cardiomyocytes (hiPSC-CMs) have emerged as a new scalable experimental model for cardiovascular research and translation^2-4^. The growth of these cells *in vitro* is being optimized for applications in regenerative medicine, drug development, cardiotoxicity testing, and other personalized medicine applications^5-7^. iPSC-CM oxygen consumption and metabolism are considered of key importance for their maturity^8-10^. For drug development and cardiotoxicity studies, human iPSC-CMs are typically grown in static culture using glass-bottom high-throughput format plates (96-well or 384-well plates). In such conditions, the only oxygen diffusion path is from the top, through the solution. Long-term studies of peri-cellular oxygen dynamics in human cardiac cells in such high-throughput plates are lacking, yet highly desirable.

Historically, oxygen sensing has evolved, starting with amperometric measurements using a Clark electrode ^11, 12^, where the electrochemical reduction of oxygen is registered by changes in electric current. This method still serves as the gold standard. Yet, it has limitations, especially for assessing oxygen dynamics in small spaces/volumes, including assessment of peri-cellular oxygen. This is due to the consumption/depletion of oxygen at the sensor during the measurement and the general difficulty in miniaturizing this type of sensor. Contactless optical methods present an alternative. A variety of fluorescent dyes have been developed to register very low (<5%) oxygen levels^13, 14^, including near-infrared indicators for *in vivo* measurements^15^. The limitations of such dyes include difficulty in providing a quantitative assessment and in covering a broader range of physiological values.

The Seahorse XF platform^16^ is a high-throughput version of an optical oxygen measurement system. It has numerous applications in rigorous metabolic profiling of mammalian cells and isolated mitochondria^17-19^, including iPSC-CMs8. It provides quantitative assessments of oxygen consumption rate (OCR) and extracellular acidification rate (ECAR), including in a 96-well format, as it offers oxygen and pH measurements through the optical sensors embedded in the tips of the fiber optics array. However, the Seahorse assay is applicable only to acute terminal measurements and therefore cannot be used for long-term tracking of peri-cellular oxygen dynamics.

Luminescence-based oxygen sensors, often combined with fiber optics, have been demonstrated to offer reliable oxygen tracking over time^20-23^. Their operation is based on dynamic oxygen quenching of fluorescence, as reflected in the Stern-Volmer relationship between oxygen concentration and fluorescence^24^. Quantitative optical oxygen sensing is typically done through life-time measurements (using frequency modulation) or through ratiometric intensity measurements with an oxygen-responsive dye (e.g. ruthenium-based) and a reference dye^20, 25, 26^). Scalability with such luminescence-based oxygen sensors has been achieved through improved matrix embedding of the dye and production of oxygen-sensing scaffolds for space-resolved measurements^27-29^, as well as through advances in visualization with high spatiotemporal resolution^30^.

In this study we characterized and validated a high-throughput platform for longitudinal optical sensing of peri-cellular oxygen in human iPSC-CMs and human cardiac fibroblasts in 96-well format within a standard cell culture incubator. The system is based on the VisiSensTD oxygen imaging system (PreSens Precision Sensing GmbH, Germany) and our high throughput microfluidics-based uninterrupted cell culture perfusion system (HT-µUPS). Results demonstrate that the system provides accurate and reproducible long-term measurements of peri-cellular oxygen levels that will be valuable for studies of cellular oxygen consumption, metabolic perturbations, and characterization of the maturation of cultured iPSC-CMs.

## Materials and Methods

### Optical oxygen sensors and peri-cellular oxygen imaging

Peri-cellular oxygen was measured using emission ratiometry. An integrated system comprised of an LED light source and an RGB camera (VisiSensTD, PreSens) imaged changes in the luminescence of optical oxygen sensors placed at the bottom of each well of a 96-well glass-bottom plate (**Fig. 1A)**. The plate was placed on top of the VisiSensTD system for continuous monitoring in a cell culture incubator. The oxygen sensor membrane (pink, **Fig. 1B**) incorporated an oxygen-responsive ruthenium dye and an oxygen-insensitive reference dye. Blue light from the LEDs positioned around the camera lens excited the two dyes. The camera imaged the entire plate to acquire oxygen-dependent changes in the luminescence ratio of the two dyes within the sensor located in each well of the plate.

**Fig 1.**
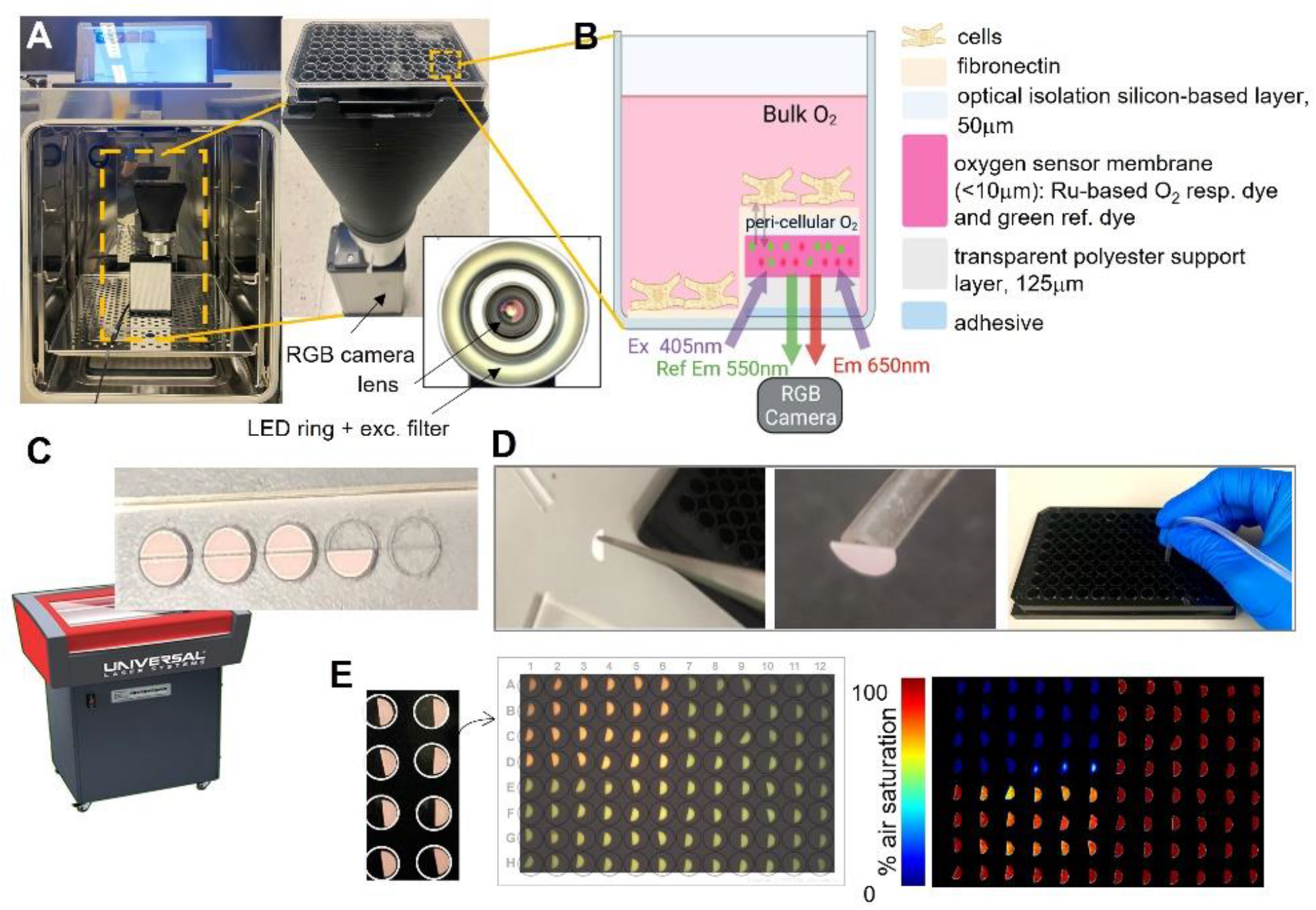
System and workflow for high-throughput optical measurements of peri-cellular oxygen. A. Incubator-deployed imaging system for 96-well plates, with an RGB camera, ring LED illuminator and bellows extension to fit a 96-well plate. B. Schematic of an optical oxygen sensor composition and placement in part of the well. Peri-cellular oxygen is imaged ratiometrically through the glass bottom of the well plate; partial coverage allows for other parameters to be measured optically, e.g. voltage, calcium etc. C. Laser-cutting of half-moon shaped oxygen sensor patches. D. Patches are detached from the adhesive backing and placed in wells using vacuum tubing. E. Images of the oxygen sensors in wells, raw optical readout and processed readings of oxygen. In the example, the upper left quadrant of the plate has been treated with oxygen-depleting Na_2_SO_3_.

The oxygen sensors were semicircles that covered half the glass bottom of each well. They were laser-cut from a larger sheet (PreSens SF-RPSu4) that consisted of an oxygen sensitive layer, a polyester support layer, and a white optical isolation layer, which was placed onto a sacrificial acrylic sheet with the adhesive facing up (**Fig. 1C)**. Semicircular sensors were then laser-cut using a 30W CO2 laser (Universal Laser Systems VLS 2.3) by placing the acrylic layer on the laser cutter bed and focusing the laser on the top of the acrylic layer. The laser cutting path was drawn in AutoCAD 2022 to cut half-moons with a radius of 2.5 mm using 15% maximum laser power and 10% maximum speed.

### Oxygen sensor attachment and plate sterilization

Oxygen sensors were attached to the bottom of the wells of 96-well plates using sterile procedures inside a laminar flow hood. Sensors were lifted with tweezers to expose the white optical blocking layer while placing a suction tube (ID < 2 mm) against this layer to hold the sensor while lowering it into a well (**Fig. 1D)**. The suction in the tube was released once the adhesive layer attached to the glass. The process was repeated to place sensors in each well (**Fig. 1E)**. Before using the plate for a cell culture experiment, the wells were sterilized with 70% ethanol inside the sterile laminar flow fume hood. After the ethanol evaporated, each well was washed three times with 1x PBS, before coating with fibronectin. After an experiment, raw images acquired by the VisiSensTD system were processed using a two-point calibration to convert the RGB values for pixels that imaged each sensor to a percentage that corresponded to the peri-cellular oxygen level. An example raw luminescence image showing the optical sensors in a 96-well plate and the processed oxygen reading is illustrated in **Fig. 1E**.

### Oxygen sensor spectral characterization

Spectra for the full range of aqueous oxygen concentration were first measured by bubbling a beaker of water with 100% O_2_ gas followed by bubbling with 100% N_2_ gas (**Fig 2A)**. A sensor attached to the bottom of the beaker was illuminated with excitation light from the VisiSenseTD system and the spectrum of emitted light was acquired once every second using a spectrometer (Ocean Optics QE-Pro, **Fig 2B)**.

**Fig 2.**
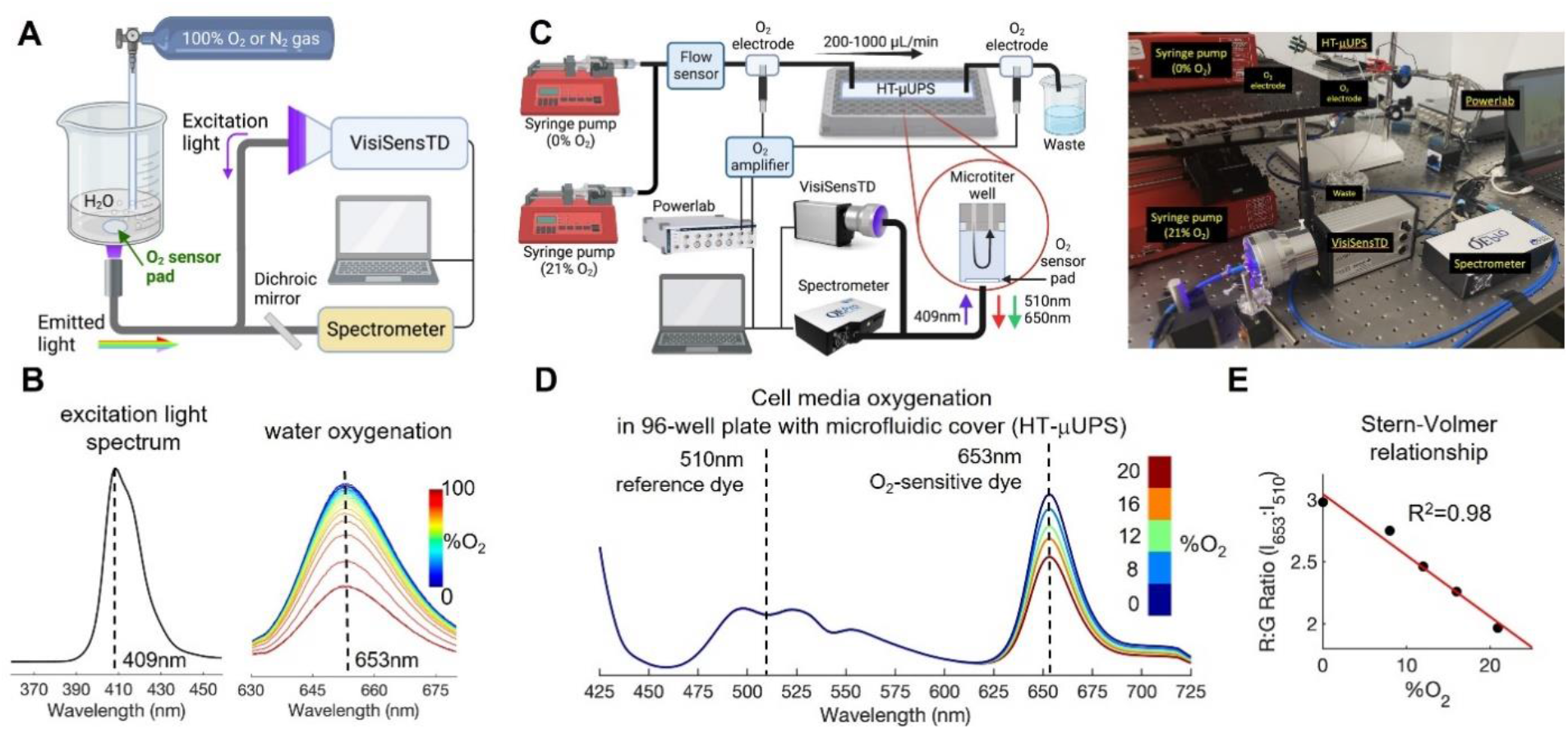
Spectral characterization of the optical system for peri-cellular oxygen imaging and confirmation of an inverse Stern-Volmer relationship. A. Schematic of the setup used to characterize oxygen-sensitive responses in a beaker of water. B. Spectral results in water: excitation light emission is at 409nm; oxygen-sensitive emission peak is at 653nm, with emission decreasing as O_2_ concentration increases. C. Schematic and photo of the setup for spectral characterization of oxygen responses in a 96-well microplate with a specialized microfluidic cover (our HT-μUPS system). The oxygen sensor pad is at the bottom of one well. D. Spectral data in cell culture media, showing oxygen-dependent spectral shifts at 653nm, consistent with results from water shown in panel B. E. An inverse Stern-Volmer relationship was derived from the ratios (intensity at 653/intensity at 510), measured from the spectra shown in panel D, which is linear for the low oxygen range considered (<20%).

Spectra for oxygen concentrations that are typical for cell cultures were acquired using a similar approach and our high throughput microfluidics-based uninterrupted perfusion system (HT-µUPS) cover for a 96-well plate ^31^ (**Fig 2C)**. Nitrogen-bubbled cell culture media (0% O_2_) and media equilibrated in room air (21% O_2_) was loaded into two separate syringes that were each placed in one of two syringe pumps (New Era 1600×2). Media from each syringe flowed through a sensor (Sensirion SLI-2000) that measured flow rate as media moved through our HT-µUPS cover to perfuse the wells of the plate (**Fig 2C)**. Flow-through Clark electrodes (Microelectrodes

Inc. MI-730) incorporated into the tubing before and after the plate measured the media oxygen concentration at those positions. Total system flow rate was maintained at 200μL/min while the flow rates of the two syringe pumps were varied to achieve mixtures of 0, 8, 12, 16, and 20 percent O_2_, which were confirmed by the Clark electrodes. The same spectrometer and fiber optic cable setup as in **Fig. 2A** was used to acquire the luminescence spectrum of a sensor within one well once every second.

Spectra collected during each characterization were plotted together to visualize changes in spectral bands corresponding to the oxygen-insensitive reference dye (centered at 510nm, “green”) and the oxygen-responsive ruthenium dye (centered at 653nm, “red”, **Fig 2D)**. Values at the peak wavelengths for red and green luminescence (**Fig 2D**, dashed lines) were used to construct the inverse Stern-Volmer relation in **Fig 2E**. For this relationship, the ratio of red to green emission was plotted against each oxygen concentration measured by the Clark electrode placed before the plate. A linear fit of ratio versus oxygen concentration was used due to the small oxygen concentration range of 0-20% that corresponds to typical cell culture conditions.

### Oxygen-impermeant HT-µUPS cover and temporal characterization

The responsiveness of the oxygen sensing system to changes in media oxygen concentration and flow rate was characterized using 96-well plates and a new oxygen-impermeant version of the HT-µUPS cover described in our previous work^31^. This cover was assembled in two components, one soft PDMS sealing gasket and one acrylic perfusion base that was CNC-milled from cast acrylic (McMaster-Carr 8560K354) and thermally bonded using a heat press (Rosineer Grip Twist) at 130°C for 3 hours. The sealing gasket was fabricated by pouring a mixture of Sylgard 184 (Dow) and Dragonskin 10 (Smooth-On) in a 1:2 ratio into an acrylic mold. Finally, an inlet and outlet were tapped (10-32 thread) for each well and fitted with stainless steel 10-32 barb-to-thread connectors (Pneumadyne).

The sensor response within the perfusion system HT-µUPS was validated using the setup in **Fig 3A**. Clark electrodes were placed before and after the cover. An optical oxygen sensor was placed at the bottom of the leftmost well of the cover. The Clark electrodes and VisiSensTD system were calibrated with water at 100% O_2_ and 0% O_2_. Water bubbled with 100% O_2_ was loaded into a syringe pump and perfused through the cover. Flow rate was measured using a flow meter (Sensirion SLI-2000) and varied between 500μL/min and 200 μL/min to determine the system’s response to changes in flow.

**Fig 3.**
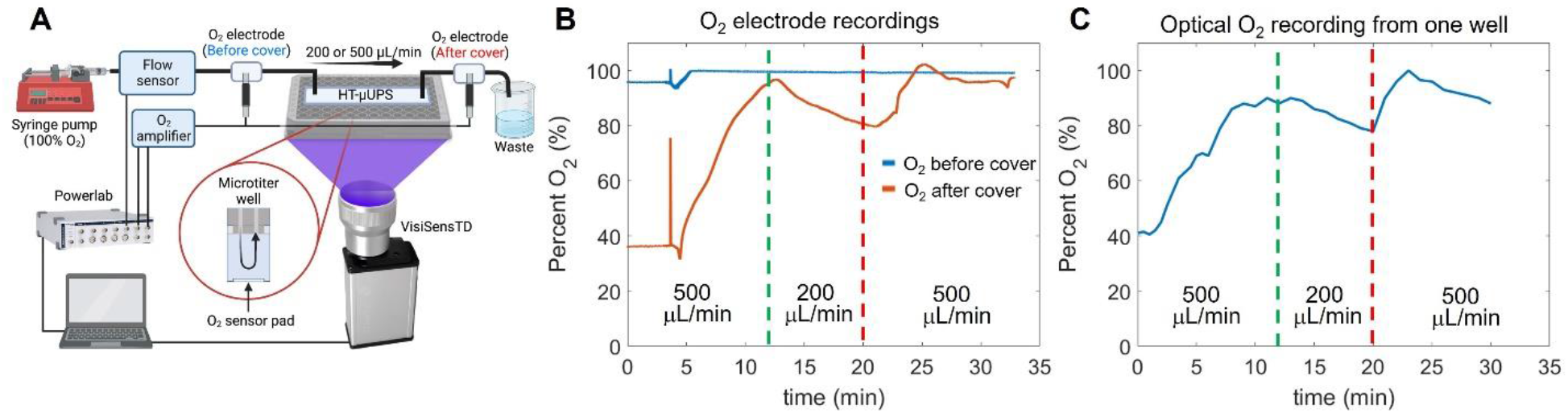
Validation of optical oxygen measurements with an electrochemical Clark electrode. A. Schematic of the setup used to perform the validation. The HT-uUPs was with an acrylic microfluidic cover. B. Measurements with the Clark electrodes positioned before and after the cover. Perfusion flow was varied to alter oxygenation of the 96-well plate (see vertical lines). C. Simultaneous measurements with the optical system reading the optical oxygen sensor within the microwell.

### Two-point oxygen calibration

A two-point calibration converted the ratio of red to green luminescence intensity imaged from the sensors into percentage of air saturation similar to ^29^. The terms ‘Cal 0’ and ‘Cal 100’ were used in system characterization experiments for 0% and 100% air saturation, and oxygen saturation levels ranging from 0% to 18.6% were displayed in cell experiments (the equilibrium oxygen partial pressure in a humidified 37°C, 5% CO_2_ incubator is 18.6%).

Two types of culture media, CDI iCell Cardiomyocytes^2^ maintenance medium (Fujifilm CDI) and cardiac fibroblasts growth medium (Cell Applications, Inc), were used. The air-saturated medium (Cal100) was medium straight from the bottle, warmed to around 37°C. And the 0% air-saturated medium (Cal 0) was medium with 5% completely dissolved Na_2_SO_3_ and placed in the 37°C water bath for 30min. In testing pH’s influence on oxygen readout, the media were adjusted with NaOH or HCL to pH 10 or pH 4.

### Cell plating and peri-cellular oxygen monitoring

Human iPSC-derived cardiomyocytes (iCell Cardiomyocytes^2^ CMC-100-012-001 from a female Caucasian donor) from Fujifilm Cellular Dynamics International (CDI), and human cardiac fibroblasts, CF (Cell Applications, Inc.) were thawed according to manufacturer’s instructions. Cells were plated (50,000 cells per well) in the wells of a 96-well glass-bottom plate containing half-moon shaped oxygen sensors, that have been sterilized and fibronectin-coated (at 50μg/ml). Culture medium exchange was done every 48 hours. In some experiments, hypoxia was induced on day five after plating by filling the wells to the top with culture medium and sealing them with oxygen-impermeable tape before readout in the VisiSense system.

The VisiSenseTD system was temperature-equilibrated in the cell culture incubator at least an hour before the start of measurements. For the whole plate oxygen monitoring in human iPSC-CMs, the peri-cellular oxygen measurements started 5 hours after the cell plating, after the switch from the cell plating medium to the cell maintenance medium. And for the hiPSC-CMs and cardiac fibroblast hypoxia comparison experiment, the oxygen monitoring started right after the hypoxic condition was established. Oxygen recordings were set to continue for 24h to 48h, with a 10 min sampling interval and 0.8 sec exposure time.

### Oxygen data analysis

Peri-cellular oxygen readings were acquired as PNG images and then analyzed through VisiSensVS software by selecting regions of interest in each image. Measured ratios were calibrated to percentage oxygen readings using two-point calibration with 5% Na_2_SO_3_ and upon saturation with ambient air. Time-dependent calibration files were applied in the first two hours of recording, considering the temperature-induced changes in transferring the plate from room temperature operation to the 37ºC humidified incubator. Whole-plate data normalization was applied by identifying the maximum and minimum ratio readouts from the whole plate throughout the whole period of recording. A scale factor was calculated by setting the maximum oxygen reading as 18.6%, the equilibrium oxygen concentration in a cell culture incubator.

## Results

### Mechanism of optical peri-cellular oxygen measurements

Optical peri-cellular oxygen measurements were based on dynamic oxygen quenching of a ruthenium dye embedded in the sensor patches on top of which cells were grown. For quantitative ratiometric readout, red and green luminescence intensity ratios were converted to percentage of oxygen saturation (pO2) using an adapted Stern-Volmer equation:

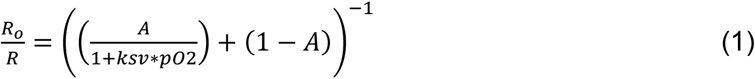

where R is the measured luminescence ratio, R_0_ is the luminescence ratio at 0% O_2_, ksv is the Stern-Volmer constant indicating efficiency of oxygen quenching, and A is 0.82, a parameter for the nonlinearity of the sensing material ^29^. The ksv was computed using the two-point calibration described in the Methods, and a time-dependent ksv was applied for the beginning of the recordings to account for temperature changes. Equation 1 was linear for the limited range of oxygenation that is typical for cell cultures (<20%, **Fig. 2E)**. However, when measured over a full range of 0-100% oxygen concentration, the inverse Stern-Volmer is expected to follow a decaying exponential function ^32^.

### High-throughput compatibility

The semicircular oxygen sensors covered half of each well in a 96-well glass bottom plate (**Fig. 1C, D**). This feature enabled multiparametric optical high-throughput measurements from the other half of each well. Such measurements include cellular action potentials, intracellular calcium transients and contractility ^33-35^. Laser cutting of the sensors was quick and reproducible (**Fig 1E**), where more than 96 half-moon sensors could be cut in less than 10 minutes. Subsequent attachment of the sensors in each well of a 96 well plate could be completed in less than one hour. Laser cutting had negligible effect on sensor performance and provided sufficient sensor area for good ratiometric measurements after selecting a region of interest from each sensor. We found that it is essential to keep the pre-cut sensors in the dark and to use them within 6 months of laser cutting for best results. Sterilization of the sensors with ethanol for cellular experiments did not affect their performance.

### System characterization and validation

The RGB images of the entire plate (1280×1024 pixels, 24bit) acquired by the VisiSensTD system provided sufficient contrast with approximately 1000 pixels per sensor, and 300 to 700 pixels per sensor region of interest (**Fig. 1E, middle)**. After calibration, the pseudocolor images of the plate clearly denoted differences in oxygenation between wells. In the example shown in **Fig. 1E, right**, the quadrants of the plate were conditioned with 0% air-saturated PBS (top left), 100% air-saturated PBS (top right), room air (bottom left), and 100% air-saturated PBS with pH=4 (bottom right). We found that solution pH had negligible influence on calibrated oxygen sensor values. The oxygen sensors also reacted differently in air than when submerged in solution.

Spectral characterization of the optical oxygen sensors and validation of the oxygen measurements with Clark electrodes confirmed that small changes in peri-cellular oxygen concentration could be accurately measured in 96-well plates (**Figs 2 & 3**). The Presens oxygen sensors (SF-RPsSU4) had a peak emission of 653 nm upon excitation with 409 nm light, and this emission decreased in response to increased oxygen concentration (**Fig. 2B**), demonstrating dynamic ruthenium fluorescence quenching by oxygen ^20, 28^. Increased fluorescence in the green band (485 – 570nm) was also measured from the sensors upon illumination with 409nm light (**Fig. 2D**), with a midpoint of 510nm. The intensity of this band did not change when oxygen concentration increased, corresponding to the oxygen insensitive reference dye that Presens has added to the sensor material. These controlled spectral measurements in a 96-well plate using our HT-µUPS system confirmed adequate spectral sensitivity of the sensors for the range of peri-cellular oxygen that is expected for cell cultures (<18.6%). The relationship of the ratio of red to green emission intensity was linear (**Fig. 2E)**, as predicted by Equation 1.

A recent study cross-validated VisiSensTD peri-cellular oxygen measurements using a fiber-optics oxygen microelectrode ^29^. In contrast, we used in-line Clark electrodes to validate measurement of peri-cellular oxygen using laser-cut semicircular sensors, the VisiSensTD system, and physiological flow rates through our HT-µUPS system in a 96-well plate (**Fig. 3A**). Our system demonstrated appropriate responsiveness to changes in media oxygen concentration and flow. During a 30min recording, media flow began at 500μL/min, was reduced to 200 μL/min, and restored to 500 μL/min (**Fig. 3A**). Oxygen values from the Clark electrode positioned at the inlet of the plate indicated that media oxygenation was near 100% and remained constant (**Fig. 3B**).

Oxygen values from the Clark electrode positioned after the plate aligned with values measured optically from the sensor at the bottom of the first well. The diameter and length of tubing and the time of the peak oxygen value (85%) measured at the bottom of the well (**Fig. 3C**, green line) corresponded to the flow rate of 500 μL/min. The drop in oxygenation at 200 μL/min indicated a loss of oxygen as the media traveled more slowly through the connection at the entrance of the HT-µUPS cover.

### Optimal exposure time

Excitation light was not uniformly distributed across the bottom of 96-well plates. This increased the spatial variance (well-to-well differences) of red:green emission ratios from the sensors for short excitation light exposure times. The dependence of sensor emission ratio variability on illumination exposure times between 0.15 to 1.5 secs was measured to identify exposure times where spatial variation was minimized (**Fig. 4A**). Spatial variation was highest for exposure times less than 0.4 sec. Spatial variation was lowest for exposure times between 0.4 and 0.9 sec. An optimal exposure time of 0.8 sec was chosen within this range as a duration for the follow up experiments. The red:green ratio for wells of a 96-well plate for an exposure time of 0.8 sec at two levels of oxygenation (0% and 100% air-saturation) and two cell culture media (hiPSC-CMs and cardiac fibroblast) varied within ± 6% (**Fig. 4B, C**), which was much lower than the typical variability between experimental groups of cultured cells.

**Fig 4.**
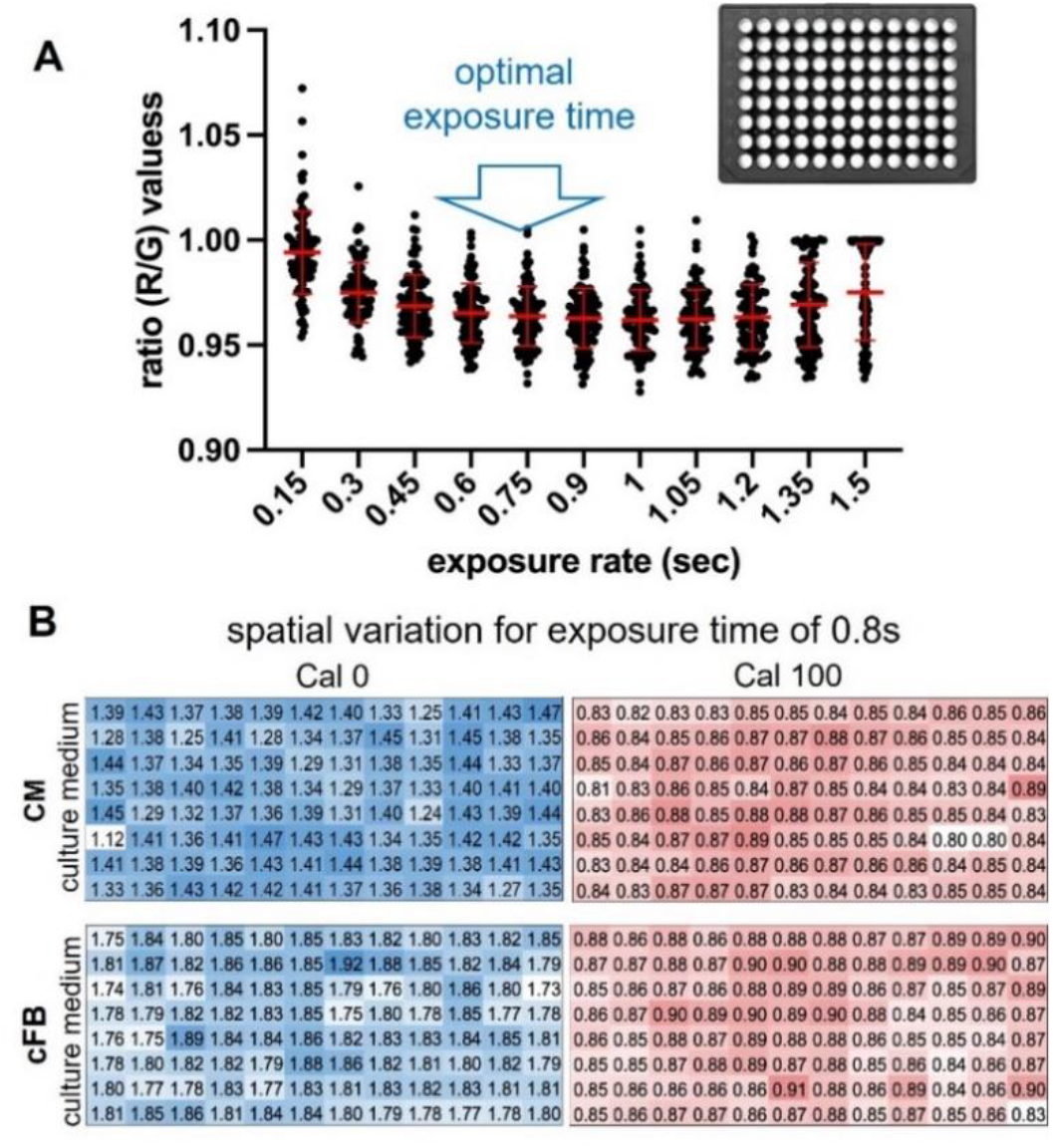
Influence of exposure time on ratio readouts and their spatial spread in 96-well plate. A. Oxygen readout (ratio values) spread across the 96-well plate as exposure time varies between 0.15 to 1.5 s. To minimize the spread, exposure was chosen in the 0.4 to 0.9 s range. B. Spatial variation across 96-well plate with 0% oxygen (Cal0), blue, and air saturated (Cal100) oxygen for culture medium used to grow human cardiac myocytes, CM (top), and human cardiac fibroblasts, cFB (bottom). 5% Na_2_SO_3_ in the medium was used for 0% oxygen calibration; exposure time set to 0.8 sec.

### Temperature effect

Temperature had a significant effect on the red:green emission ratio for each sensor, which was evident after placing a 96-well plate at room temperature in the incubator maintained at 37ºC. The effect of temperature on emission ratio, as a plate was warmed to 37ºC, was measured over 4 hours after placing a plate in the incubator (**Fig. 5A**). Wells contained media for hiPSC-CMs or media for cardiac fibroblasts, and wells had an oxygen level of either 0% or 100% air-saturation. Oxygen images were acquired every 5 minutes. The average red:green ratio for each combination of media and oxygenation changed over the first hour (**Fig. 5A)**. Ratios for wells having media at 100% air-saturation were similar throughout the 4 hours and changed approximately 10% in the first hour. Ratios for wells having media at 0% changed as much as 60% in the first two hours and the change was greater for wells with fibroblast media. This result was likely due to differences in the content of the two types of media. The Stern-Volmer ksv values (Equation 1) computed from the ratios for wells having media at 0% also varied within the first hour (**Fig. 5A**, inset). These results necessitated the development of a protocol to correct for the effect of changes in well temperature as a plate was warmed to 37ºC and for the effect of cell culture media content that determines media color.

**Fig 5.**
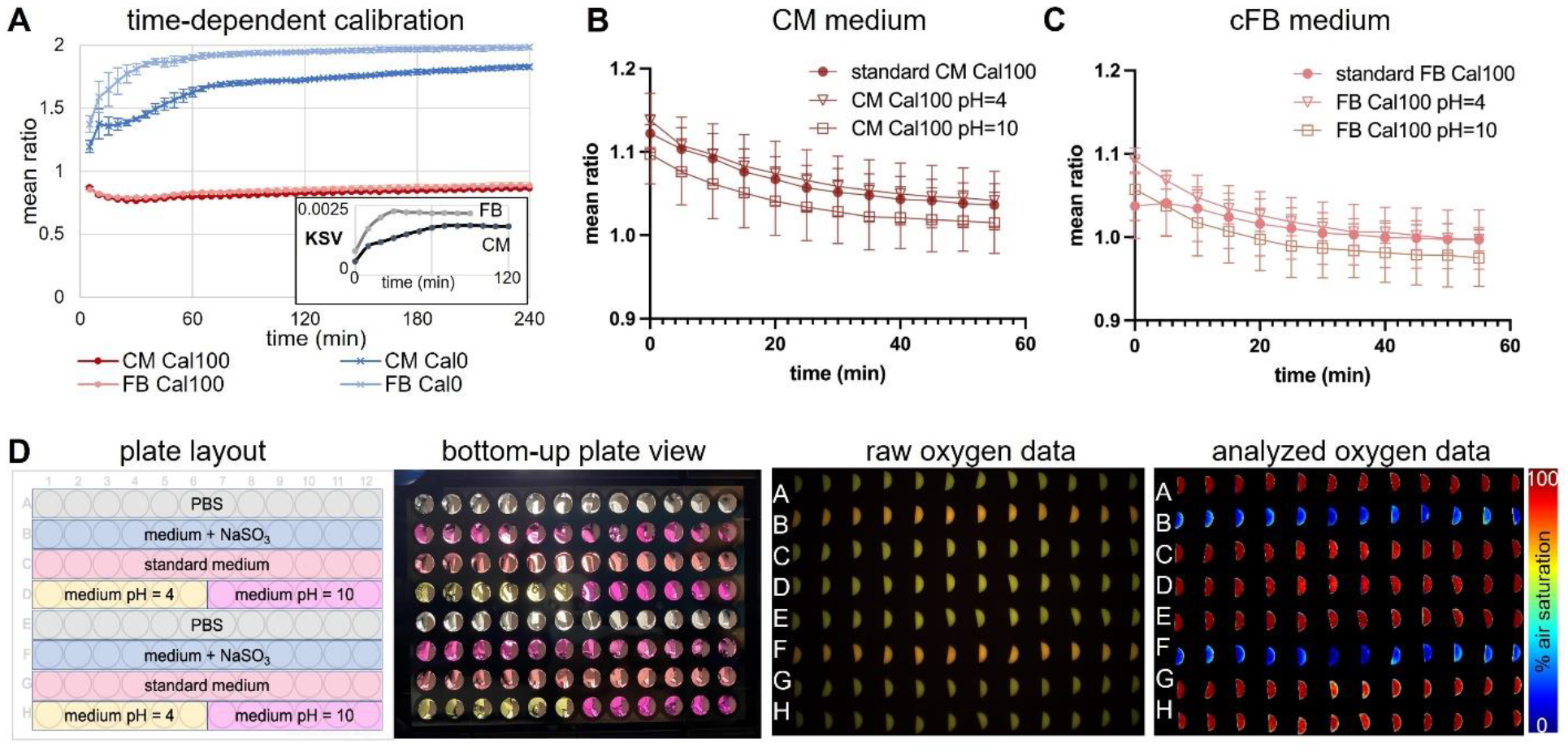
Time-dependent calibration, effects of pH and medium color on the oxygen readings. A. Time-dependent calibration over 4 hours, done in culture medium for human cFBs and human iPSC-CMs in a 37ºC cell culture incubator, n=96 wells. Over the first 40min, there is a temperature-dependent rise in the ratio reading for the Cal0 values. Inset shows that the derived Stern-Volmer coefficient, KSV, varies in this initial period for both culture media. B. Time-dependent calibration changes of CM medium with different pH = 4, 7.4 (standard) and 10. Oxygen readouts were continuously recorded for an hour in 5 min intervals. The mean ratio of air-saturated CM medium with standard and altered pH was plotted over time (mean±SE, n=12-24 samples). C. Same as B, but for cFB culture medium. D. Full 96-well plate characterization of medium color and pH’s influence on oxygen reading. Conditions are listed in the plate layout. Although pH visibly altered culture medium color (bottom-up view), the analyzed oxygen data were not influenced by medium color or pH difference.

### Effects of media color and pH

The substrate, salt, and chemical content of the hiPSC-CM media and the cardiac fibroblast media was different, resulting in each culture media having a distinct color. Media pH also determines media color and changes in pH could have an independent effect on sensor luminescence. The effect of media color and pH on sensor emission ratio was studied using 96-well plates and the VisiSensTD system. Wells contained either PBS, hiPSC-CM media, or cardiac fibroblast media and the pH of each well was set to be either 4 or 10 (**Fig. 5B**, plate layout). Differences in the color of the media in each well was clearly visible (**Fig. 5B**, bottom-up plate view). However, color differences were not evident in images of the emission ratio (**Fig. 5B**, raw oxygen data). Color differences were also not evident in pseudocolor images after computing the percent oxygen concentration of each well using the emission ratios (**Fig. 5B**, analyzed oxygen data). To determine if pH had a direct effect on sensor luminescence, emission ratio of each media having 100% air-saturation at standard pH of 7 or a pH of 4 or 10 was measured once every 5 minutes for one hour using the VisiSensTD system (**Fig. 5C, D**). No significant difference was detected between media having a pH of 7 and 4. The emission ratio was consistently lower for media having a pH of 10, which is highly alkaline and has less biological relevance for cell culture experiments.

### Long-term monitoring of cell culture peri-cellular oxygen concentration

After validation, characterization, and developing a robust calibration protocol, the VisiSensTD oxygen imaging system was used to continuously measure peri-cellular oxygen concentration from cardiac cells cultured in a 96-well plate. Studies were conducted for time intervals ranging from 24 to 48 hours (**Fig. 6**). In one experiment, peri-cellular oxygen was measured every 10 minutes over 48 hours from hiPSC-CMs plated in every other row of a 96-well plate (**Fig. 6A**). Two control wells had oxygen sensors and media but no cells. Peri-cellular oxygen dropped dramatically to <5% within 10 hours in all wells that had cells. Oxygen concentration in control wells was constant at 18.6%, corresponding to equilibration of the media with the incubator oxygen concentration. In wells having cells, two patterns of peri-cellular oxygen dynamics emerged after 10 hours (**Fig. 6A, lower left**): one of monotonic oxygen depletion and another of intermittent depletion. In cases of monotonic depletion, after the initial oxygen drop to <5%, peri-cellular oxygen monotonically and slowly decreased to a steady-state level. In intermittent depletion, peri-cellular oxygen oscillated intermittently after the initial oxygen drop. For the results shown in **Fig. 6A**, monotonic depletion was observed in 11 out of 38 wells (29%) and intermittent depletion occurred in 27 out of 38 wells (71%). Oscillatory intermittent depletion occurred in different wells within the rows of a plate, confirming that oxygen dynamics were independent of the position of a well within a plate.

**Fig 6.**
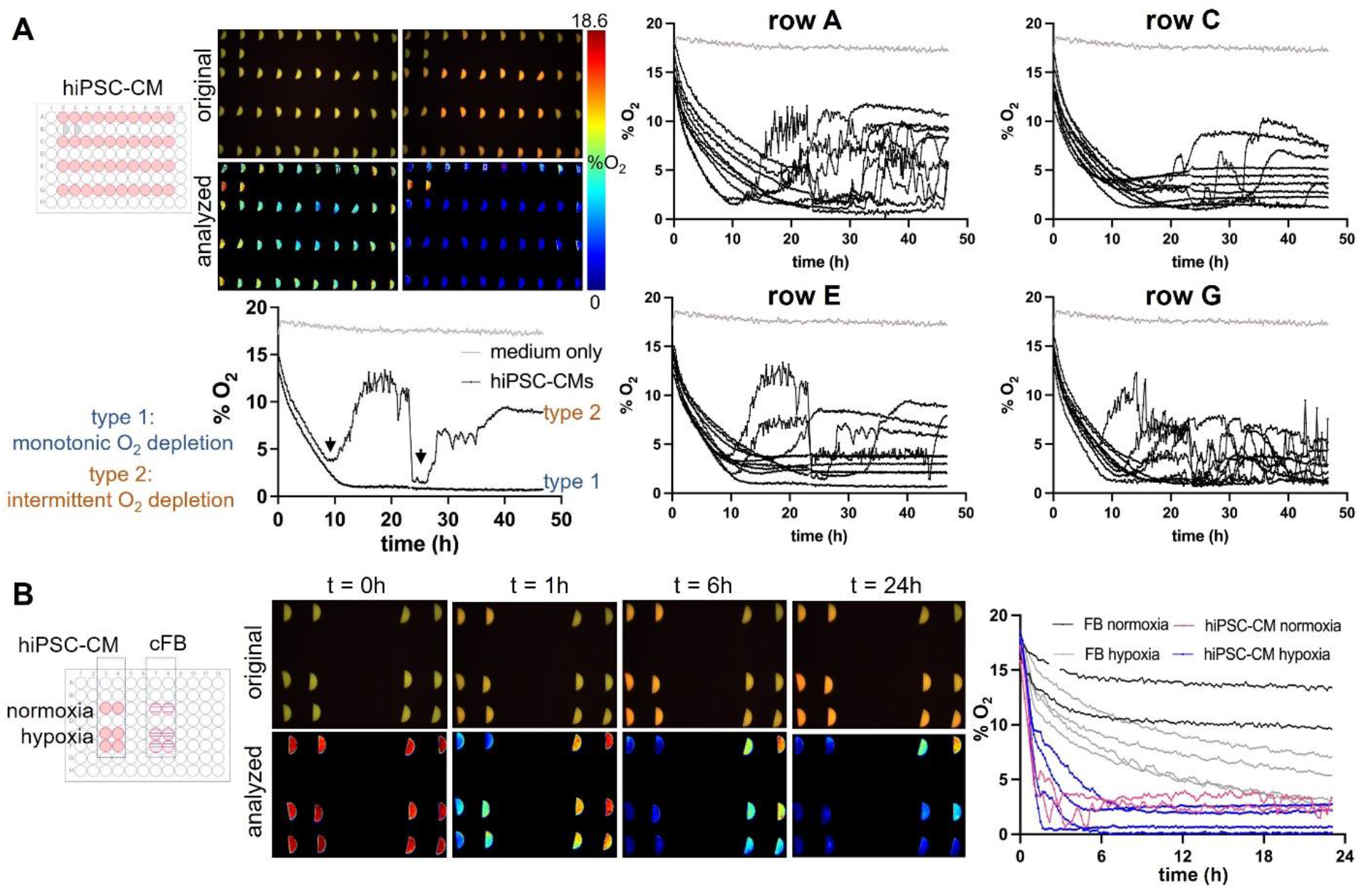
Optical measurements of peri-cellular oxygen in human iPSC-cardiomyocytes and human cardiac fibroblasts, cFBs. A. Continuous optical sensing of peri-cellular O_2_ in human iPSC-CMs over 48h. Peri-cellular oxygen levels drop to <5% in <10h for most samples. Two types of hypoxia responses are seen: (1) monotonic O_2_ depletion to a steady-state, and (2) intermittent O_2_ dynamics after an O_2_ dip below 5% (black arrows). Type 1 was observed in about 30% of the samples in this plate, while type 2 was observed in the remaining 70% of the measured n=40 wells. 18.6% oxygenation corresponds to the maximum (100%) oxygen saturation in a CO_2_ incubator. B. Example records of hiPSC-CMs and cFB peri-cellular O_2_ over 24h, under normoxic and hypoxic conditions. Peri-cellular O_2_ in cFBs decreases slower and stabilizes to a higher level; induced hypoxia (filling the wells with solution to the top and sealing them) sped up O_2_ the depletion in cFB. The hiPSC-CMs showed rapid depletion of O_2_ within 6h with faster initial phase of drop in the sealed samples, which then bounced back to slightly higher steady-state values.

Oxygen dynamic patterns were studied in cardiac fibroblast and hiPSC-CM cultures over 24 hours to determine if depletion occurs more rapidly in cells having higher metabolic rate (**Fig. 6B)**. Depletion was hypothesized to occur more rapidly in hiPSC-CM cultures due to the higher oxygen consumption of contracting cells. The two cell types were cultured in separate sets of 6 wells of the same plate. Two of those wells were filled with the typical 200μL of media to provide normoxia while the other four wells were filled with 300μL of media and sealed with oxygen-impermeable tape to generate hypoxia (**Fig. 6B, left)**. Peri-cellular oxygen was depleted within 6 hours for normoxic and hypoxic cultures of hiPSC-CMs. Oxygen depletion was much slower for fibroblasts, where over 24 hours oxygen did not drop below 10% for normoxic cultures and most hypoxic cultures maintained an oxygen level above 5% (**Fig. 6B, right)**. These results confirm the hypothesis and also demonstrate the utility of optical oxygen sensing in providing long-term measurements that reveal cell-type differences in the baseline and fluctuations of peri-cellular oxygen concentration.

## Discussion

Oxygen consumption and defense mechanisms against hypo-/hyperoxia are critical to sustaining life, as O_2_ is a key component of energy (ATP) production in the mitochondria. However, when in excess, oxygen is inherently toxic due to its chemical reactivity and the generation of reactive oxygen species (ROS)^1^. Although the complexity of measuring peri-cellular oxygen levels in cell culture has been recognized since at least 1970^1^, doing so is not routine in cell culture studies. In fact, peri-cellular oxygen measurements are rare, and no long-term measurements are available from cultures of human iPSC-CMs. A major obstacle has been the lack of user-friendly monitoring systems and reliable non-invasive methods for long-term oxygen monitoring in common cell culture formats such as 96-well plates.

In this study, the Presens ratiometric optical oxygen sensors and camera-based imaging system were adopted to track peri-cellular oxygen dynamics in human cardiac cells cultured in a 96-well plate. We developed a scalable technique for reproducible cutting and positioning of semicircular optical oxygen sensors into glass-bottom HT plates (**Fig. 1**), to cover half of each well, leaving the other half for multimodal structural and functional imaging. We validated the optical oxygen readings with traditional Clark electrodes positioned in line in bulk measurements and during perfusion within 96 wells using our specialized microfluidics-based cover. The spectral, spatial, and temporal properties of the system were systematically characterized (**Figs. 2-4**). Optimization of the exposure time and the two-point calibration (**Figs. 4-5**) allowed continuous oxygen measurements in a standard incubator that were robust and reliable.

Our results are in agreement with *in vivo* estimates of peri-cellular oxygen levels in the heart, where oxygen tension in the immediate vicinity of the cardiomyocytes is in the range of 3-6%1. Namely, for human iPSC-CMs, in the static culture conditions of a glass-bottom 96-well plate, we observed a drop to < 5% oxygen within several hours of media exchange. After this initial drop, the cells either settled to a steady state of low pericellular oxygen, likely due to the balance between active oxygen consumption and passive diffusion, or exhibited an intermittent oxygen dynamics (**Fig. 6)**. These two types of responses are interesting novel observations and require further investigation. In contrast to the cardiomyocytes, the non-contracting cardiac fibroblasts exhibited higher peri-cellular oxygen levels after dropping to a stable steady-state, as expected for cells with lower oxygen demand.

*In vivo*, cardiac cell normoxia is maintained by feedback mechanisms that tightly regulate coronary blood^36^ and by oxygen-on-demand provided by the excellent oxygen carrying capacity of hemoglobin. *Ex vivo*, in the absence of adequate oxygen buffering, perfused working hearts experience hypoxic conditions^37, 38^. *In vitro*, cardiomyocytes may experience hyperoxic, normoxic or hypoxic conditions1 depending on their density, electromechanical activity, mass transport conditions and the shortest path to the ambient oxygen supply. For human iPSC-CMs, oxygen tension was recognized in early work as a key variable for optimizing cellular differentiation and maturation; transient control of oxygen level is used during the differentiation of iPS cells into cardiomyocytes^39, 40^. Hypoxia signaling is also a foundational physiologic component of mature cardiomyocytes, where it intimately regulates electromechanical function, including ion channel current and protein expression^1^, ^41, 42^. Based on this, longitudinal label-free monitoring of peri-cellular oxygen, in a high-throughput manner, provides unique insights into the metabolic state of the cells, and potentially can be correlated with their level of maturation.

The system described here is best suited for imaging the oxygen concentration of two-dimensional multi-cell structures, such as monolayers. This is a limitation, considering the growing popularity of three-dimensional cell constructs, including cell spheroids and microtissues. A potential way to extend the described label-free oxygen sensing approach to 3D structures is inspired by recent work of others, where optical sensors have been mounted on transparent prisms^29^ and oxygen-sensing cell culture vessels have been thermoform-molded to line the wells of spheroidal plates^43^. Future work includes automating the placement of sensors into 96-well plates using the scalable technique described and manually demonstrated here. Coupling label-free measurements of peri-cellular oxygen with label-free measurements of cardiac electromechanical waves35 will also provide valuable insights into the interplay between cellular activity and oxygenation state.

## Data Availability

The data underlying this study are available in the published article.

## Acknowledgements

This work was supported in part by grants from the NIH-NHLBI R01HL144157 and from the NSF EFMA 1830941.

